# Immunity against Mycobacterium avium induced by DAR-901 and BCG

**DOI:** 10.1101/2024.09.03.611096

**Authors:** Getahun Abate, Krystal Meza, Chase Colbert, Christopher Eickhoff

**Affiliations:** Division of Infectious Diseases, Department of Internal Medicine, Saint Louis University, Saint Louis, Missouri, USA

**Keywords:** *M. avium*, BCG, DAR-901, immunity, BALB/c

## Abstract

The prevalence of pulmonary nontuberculous mycobacteria (NTM) is increasing in Europe and North America. Most pulmonary NTM are caused by *Mycobacterium avium* complex (MAC). The treatment of pulmonary MAC is suboptimal with failure rates ranging from 30% to 40% and there is a need to develop new vaccines. In this study, we tested the ability of two whole cell vaccines, DAR-901 (heat killed *M. obuense*) and BCG (live attenuated *M. bovis*), to induce MAC cross-reactive immunity by first immunizing BALB/c mice and then performing IFN-γ ELISPOT assay after overnight stimulation of splenocytes with live MAC. To study the ability of these vaccines to protect against MAC infection, BALB/c mice were vaccinated with DAR-901 (intradermal) or BCG (subcutaneous or intranasal) and challenged with aerosolized MAC 4 weeks later. Some mice vaccinated with BCG were treated with clarithromycin via gavage. Lung CFU in immunized mice and unvaccinated controls were quantified 4 weeks after infection. Our results showed that i) DAR-901 induced cross-reactive immunity to MAC and the level of MAC cross-reactive immunity was similar to the level of immunity induced by BCG, ii) DAR-901 and BCG protect against aerosol MAC, iii) mucosal BCG vaccination provided the best protection against MAC challenge, and iv) BCG vaccination did not interfere with anti-MAC activities of clarithromycin.

## Introduction

Nontuberculous mycobacteria (NTM) are all mycobacteria except *Mycobacterium tuberculosis* (Mtb) and *M. leprae*. There are more than 180 NTM species but only a few are clinically relevant [1]. NTM can affect any organ in the body, but the pulmonary form is the most common form in patients who are negative for human immunodeficiency virus (HIV) [2]. In the US and Western Europe, the prevalence of pulmonary NTM in HIV-negative patients has been increasing over the last decade [2-8]. The reasons for these recent increases in the prevalence of pulmonary NTM is not clearly known but advances in extending the life expectancy of patients with underlying chronic lung diseases such as cystic fibrosis and COPD, and wide application of immunosuppressive medications in medicine may have played a role [9-13].

In North America and parts of Europe, *M. avium* complex (MAC) is the most common NTM species associated with pulmonary NTM [8, 14]. Regardless of the causative NTM species, management of pulmonary NTM is extremely difficult. Treatment requires the use of multiple drugs for at least 12 months from the first culture-negative results which in most cases means more than 18 months of treatment [15]. Despite treatment for several months, failure rates in the range of 30%-40% have been reported [14, 16, 17]. Thus, there is an urgent need to develop new vaccines and new therapeutics. This urgency was emphasized in the last NTM workshop organized by the National Institute of Allergy and Infectious Diseases [18].

In the last decade significant advances have been made in understanding tuberculosis (TB) immunology and bringing new vaccines as well as immunotherapeutics to clinical trials. Lessons learned in the TB field could be very useful for work on pulmonary MAC. Like Mtb, MAC is an intracellular pathogen, and its control relies primarily on mounting effective cell-mediated immunity [19, 20]. This study was performed with the objective of testing the ability of DAR-901 and bacillus Calmette Guerin (BCG) to induce MAC immunity and evaluate their potential for prevention of pulmonary MAC in a murine model.

## Materials and Methods

### Reagents and Bacterial culture

MAC *(M. avium*, ATCC 700898) and Tice bacillus Calmette-Guerin (BCG, Merck) were used. Large bacterial lots of MAC and BCG were grown in roller bottles at 37 °C in albumin-dextrose-catalase (ADC)-supplemented Middlebrook 7H9 broth, aliquoted in 1-ml volume and stored at -80°C.

DAR-901, a vaccine that contains heat-inactivated *Mycobacterium obuense*, was used in some assays. DAR-901 in 2 mL vials containing 0.3–0.4 mL of a 1 mg/mL suspension of heat-inactivated organisms was kindly provided by Dr. Fordham von Reyn (Geisel School of Medicine at Dartmouth, Hanover, New Hampshire).

### Animals

Animal experiments were reviewed and approved by the Saint Louis University Institutional Animal Care and Use Committee (IACUC). Animal experiments were performed with 6-to 8-week-old female BALB/c mice as described previously [22].

### Vaccination and routes of administration

BCG was administered intranasally (IN), subcutaneously (SC) or intradermally (ID). Aliquots of BCG were thawed and pelleted by centrifuging at 3700 rpm for 15 minutes at 4°C. Pellets were resuspended in phosphate buffer saline (PBS). Mice that received IN BCG had 1 × 10^7^ BCG delivered in 40 µL doses split between nostrils. Groups of mice that received SC BCG received 1 × 10^7^ bacteria in 100 µL to base of tail. Mice that received ID BCG had 1 × 10^7^ in 50 µl PBS delivered to the left side of shaved abdomens. DAR-901 was diluted to various doses (0.1 to 2.5 mg/dose) in PBS and administered in a total volume of 50 µL using insulin syringes (BD Biosciences, San Jose, CA). Mice were anesthetized with ketamine/xylazine cocktail intraperitoneally before IN or ID vaccination.

### Quantifying lung colony forming units (cfu) after MAC infection

Mice were infected with aerosolized MAC. A total of 3 -7 mice were sacrificed on day 0 and at weeks 2,4, 6, and 8 post infection. Lungs were homogenized in sterile PBS using a bead mill homogenate, serially diluted and plated in duplicate for CFU quantification on 7H11 agar media. For mice vaccinated with BCG, additional oleic-albumin-dextrose-catalase (OADC)-supplemented Middlebrook 7H10 agar media containing isoniazid at a concentration of 1 µg/ml was used to inhibit the growth of BCG. Plates cultured at 37°C were read every week and the CFU counts were finalized on week 4.

### Measuring recruitment of T cells to the lungs following BCG vaccination

At different time points post vaccination, mice vaccinated with BCG via SC or IN route were injected with fluorescently labeled anti-CD45 intravenously (i.v.) and then euthanized 3-5 minutes later, allowing sufficient time to stain cells in the vasculature but not those in the tissues. Lungs were extracted and digested with collagenase/DNase. Then, single cell suspensions were prepared for flow cytometric studies. Absolute numbers of CD3+CD44+CXCR3+ were calculated by multiplying the total number of viable cells recovered based on trypan blue staining and the percentage of the specific T cell subset detected by flow-cytometric analysis.

### MAC challenge

Two-three weeks before mice experiments, aliquots of MAC (ATCC 700898) were thawed and cultured in fresh ADC-supplemented 7H9 media without Tween. On the day of challenge, mycobacterial suspensions were centrifuged, and pellets resuspended in PBS. The optical density (OD) of suspension was measured at 600 nm. The suspension was diluted to adjust optical density to OD of 0.7. We estimated 3.1 × 10^7^ CFU per OD unit based on results from previous titration experiments. MAC at an estimated final concentration of 2 ×10^7^ CFU/ml was added to the nebulizer and delivered via the aerosol route using Glas-Col Inhalation Exposure System (IES) (Glas-Col Inc., Terre Haute, IN, USA). Some animals were euthanized immediately post-exposure to quantitate the delivery dose using methods described above.

To study the effects of BCG vaccination on anti-MAC activities of clarithromycin, BCG-vaccinated and unvaccinated mice were infected with aerosolized MAC. Two weeks after infection, some mice were treated with clarithromycin at concentrations ranging from 6.25 to 100 mg/kg for 4 weeks before they were euthanized. Clarithromycin was administered in a 0.2 ml volume by esophageal cannula (gavage) 5 days per week. Percent inhibition was calculated as follows: % inhibition = 100 − [100 × (CFU in mice with intervention/mean CFU in mice without intervention]. Intervention includes BCG vaccination and/or clarithromycin treatment.

### Measuring vaccine-induced MAC immunity

We used IFN-γ ELISPOT assay to measure MAC-specific T cell immunity. The assay was done as described previously [23]. Briefly, mice were euthanized at different timepoints and splenocytes (5 × 10^5^ cells/well) were stimulated overnight with live BCG at a multiplicity of infection (MOI) of 3, *M. avium* at MOI of 3, or DAR-901 at 2 and 10 µg/ml. Cells rested in media alone were used as negative controls. After overnight incubation at 37 °C, ELISPOT plates were developed using biotinylated anti-IFN-γ (BD Biosciences clone XMG1.2), streptavidin horseradish peroxidase (SA-HRP; Jackson ImmunoResearch), and AEC development solution as per manufacturer recommendations (BD Biosciences). IFN-γ producing spots in each well were enumerated using a C.T.L. ImmunoSpot analyzer and software. The results are presented as spot forming cells (SFC, mean ± SE) per million splenocytes.

## Results

### DAR-901 and BCG induce MAC cross-reactive immunity

DAR-901, like BCG, is a whole cell mycobacterium. Unlike BCG, DAR-901 contains a killed NTM. We used the intradermal route for administration of DAR-901 because that was the route used in animal and human studies for TB immunity or protection [24, 25]. Similarly, we tested 2 and 3 doses of DAR-901 vaccination since multiple doses were required in previous works to enhance TB immunity [21, 24, 25]. In our initial experiments, we compared 2 and 3 injections of three different concentrations of DAR-901 ranging from 0.3 mg to 2mg. Figure 1A show that both 2 and 3 doses of DAR-901 induce robust MAC-specific T cell immunity. Subsequently, we compared MAC cross-reactive immunity induced by DAR-901 with immunity induced by BCG administered via the same route. Figure 1B shows that DAR-901 induces the same level of MAC immunity as BCG (difference not significant).

**Figure 1.**
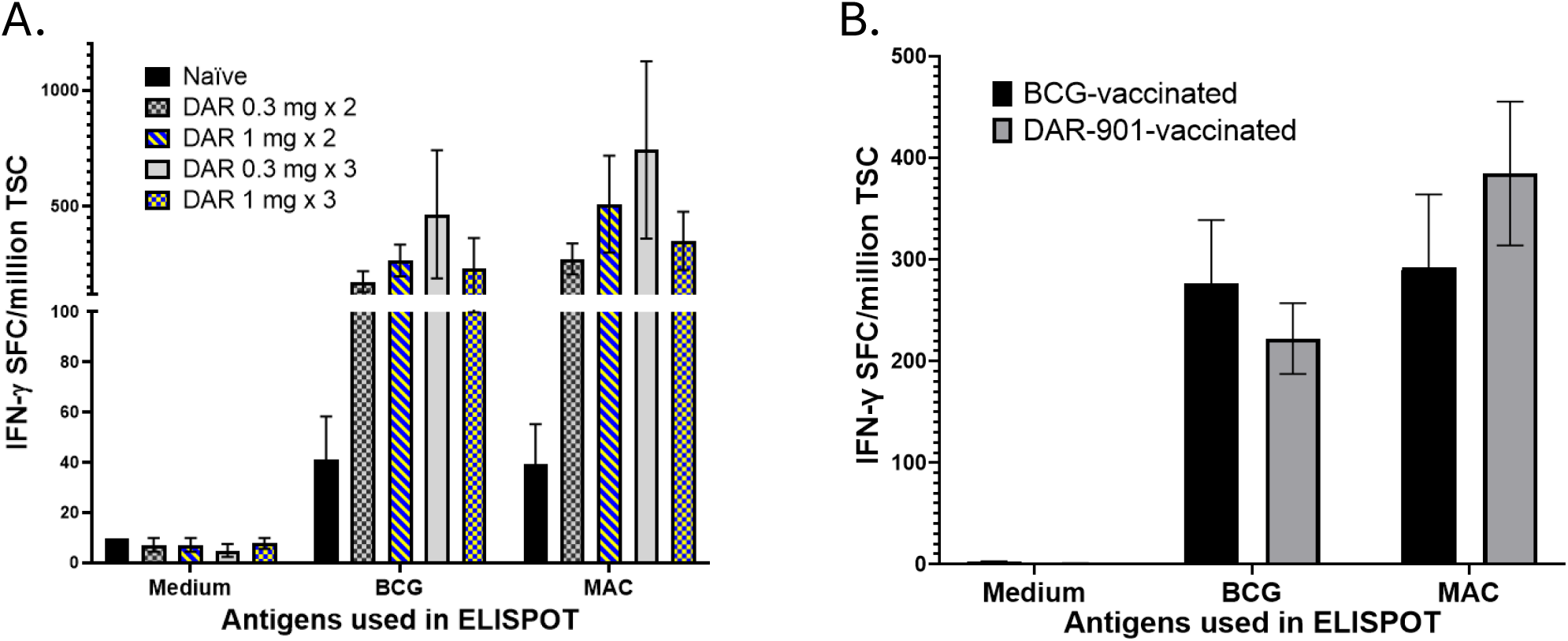
*M. avium* cross reactive immunity induced by vaccination with BCG and DAR-901. Six-to 8-week-old female BALB/c mice were vaccinated with BCG (1 × 10^7^ intradermal) or different concentrations (0.3 mg, 1 mg and 2 mg) of DAR-901 (2 or 3 intradermal doses a week apart). Four weeks after last vaccination, mice were euthanized and splenocytes were used for IFN-γ ELISPOT assay. In the ELISPOT assay, splenocytes (5 × 10^5^ cells/well) were stimulated overnight with live BCG at multiplicity of infection (MOI) of 3, *M. avium* at MOI of 3, or media alone as a negative control. IFN-γ producing spots in each well were enumerated using a C.T.L. ImmunoSpot analyzer and software. The results are presented as spot forming cells (SFC, mean ± SE) per million total splenic cells (TSC). (A) shows results from mice (4 mice /per group) vaccinated with BCG or 2 and 3 dose of 0.3 mg or 1 mg DAR-901 a week apart. Both BCG and different doses of DAR-901 induced MAC cross-reactive immunity. (B) shows results from experiments on larger number of mice (8 mice per group) vaccinated with BCG (ID), 2 doses of 0.3 mg DAR-901 a week apart or left unvaccinated. DAR-901 (2 doses of 0.3 mg) induced BCG-reactive and MAC-reactive immunity similar to immunity induced by BCG. MAC cross-reactive immunity induced by MAC was significantly higher than the unvaccinated group (P< 0.01, ANOVA).

### DAR-901 and BCG protect against MAC aerosol challenge in mice

To study the relevance of vaccine-induced immunity for MAC protection, we optimized a murine model. After low dose aerosol infection, MAC grows and reaches its peak after 4 weeks (Figure 2). The number of cfu in the lungs on week 4 was about 10 times higher with mice-adapted MAC (i.e., MAC passaged in mice lung at least once) (data not shown). Therefore, in subsequent experiments, we used mouse-adapted MAC. Figure 3 shows DAR-901 and BCG (both systemic and mucosal BCG) vaccinations protect against aerosol challenge (p< 0.001). Interestingly, mucosal BCG vaccination provided the best protection. BCG vaccination, systemic and mucosal, leads to an increase in recruitment of T cells to the lungs. Lung T cell recruitment reached its peak 10 days post vaccination. In mice with SC BCG vaccination, the number of T cells recruited to the lungs decline on week 28 but remained high in mice vaccinated with IN BCG (Figure 4).

**Figure 2.**
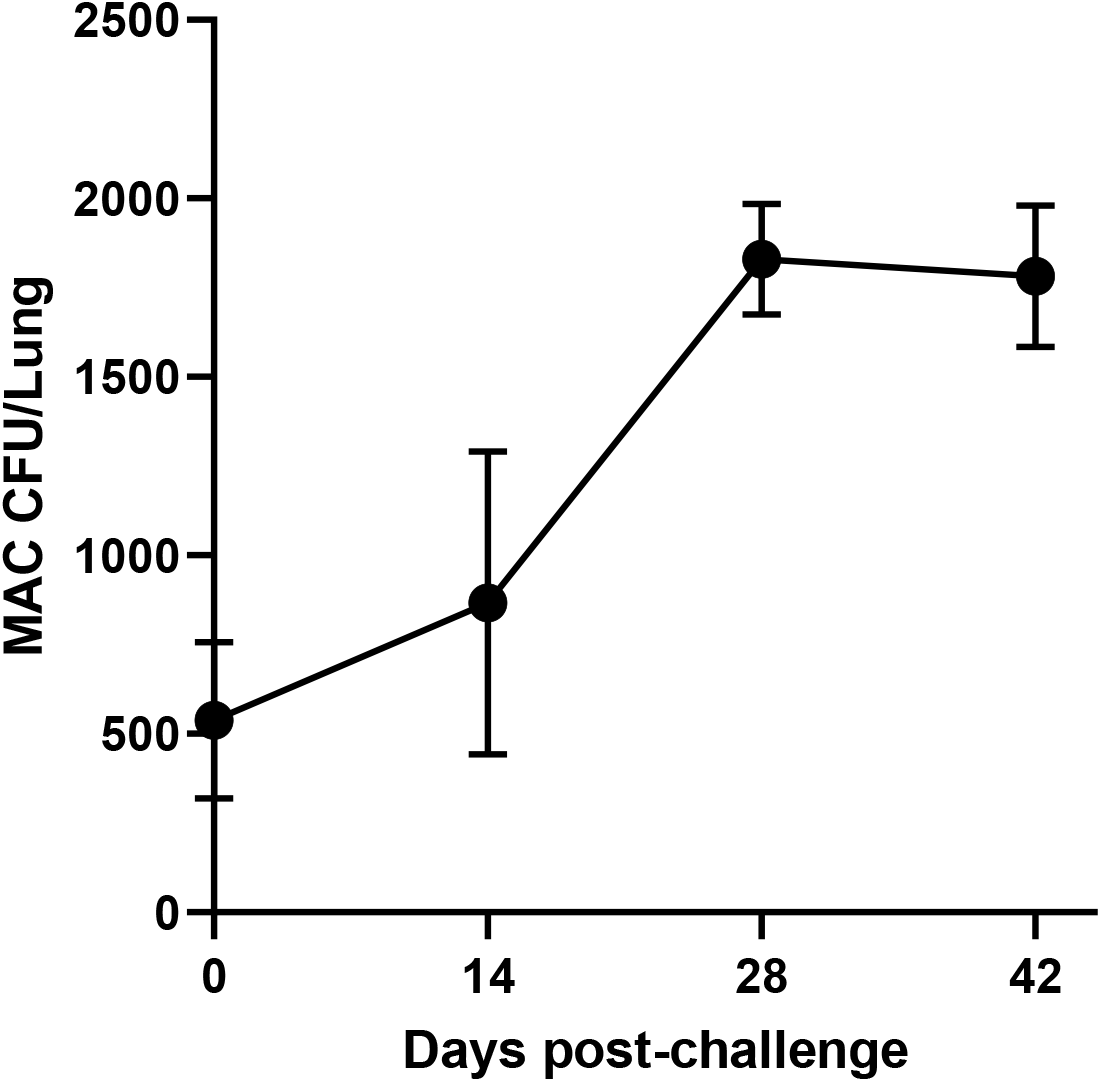
Effects of BCG vaccinations on the growth of *M. avium* in lungs. A). Mice were infected with aerosolized *M. avium* at concentration of 2 ×10^7^ CFU/ml and euthanized on day 0, 14, 28 or 42. Homogenized lungs were cultured on 7H10 media. M. avium reaches peak growth 4 weeks after challenge.

**Figure 3.**
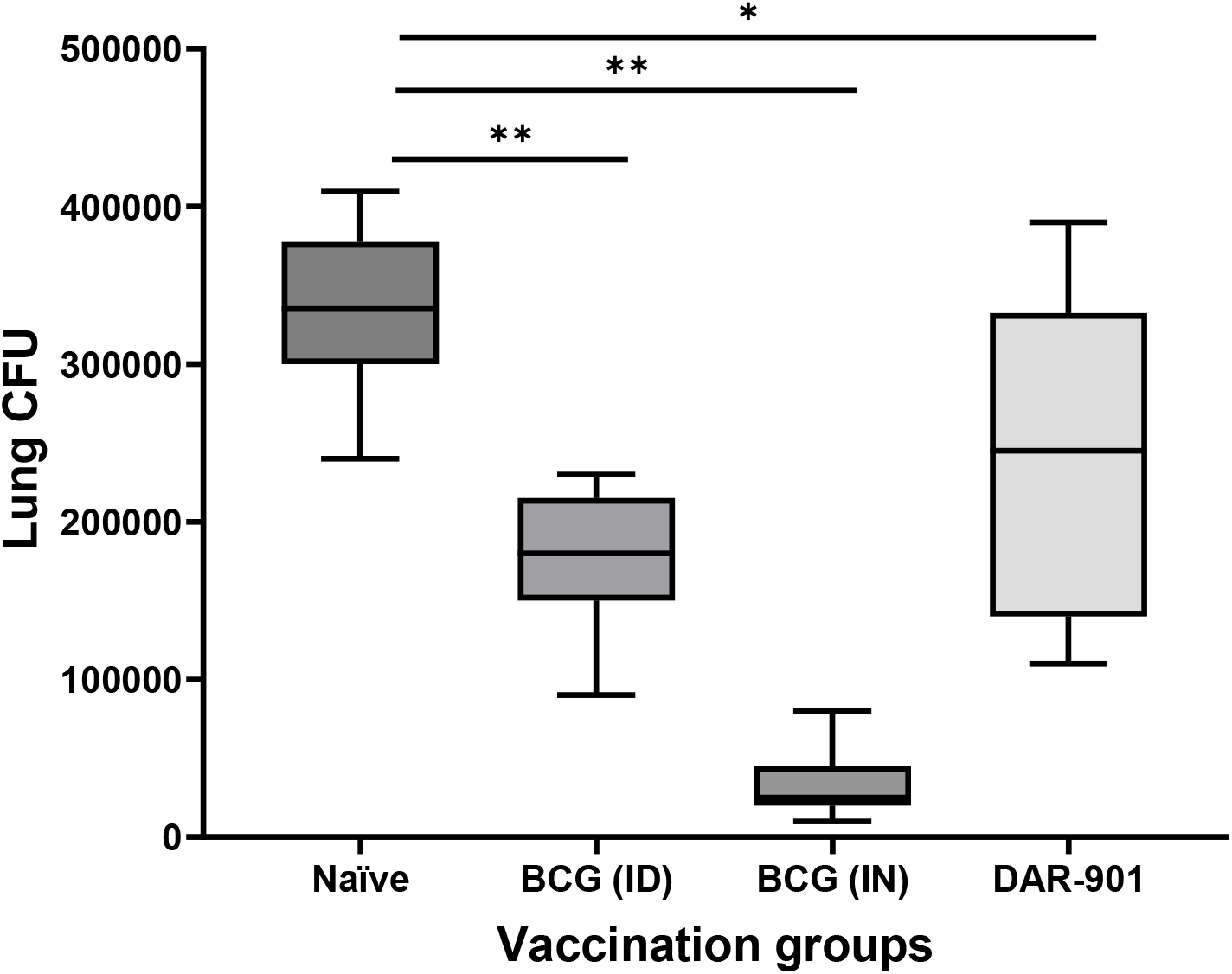
Effects of vaccinations on the growth of M. avium in lungs. Female BALB/c mice were vaccinated with BCG ID (1 × 10^7^ bacteria in 100 µL to base of tail), BCG IN (1 × 10^7^ CFU total in both nostrils) or DAR-901 (2 doses of 0.3 mg ID, 1 week apart). Four weeks after last vaccination, mice were infected with aerosolized MAC at a concentration of 2 ×10^7^ CFU/ml. All mice were euthanized 4 weeks after infection and MAC lung CFU quantified by culturing on 7H10 media. Lung CFU was significantly lower in mice vaccinated with ID BCG (P<0.001), IN BCG (p<0.001) and two doses of 0.3 mg DAR-901 (P <0.01).

**Figure 4.**
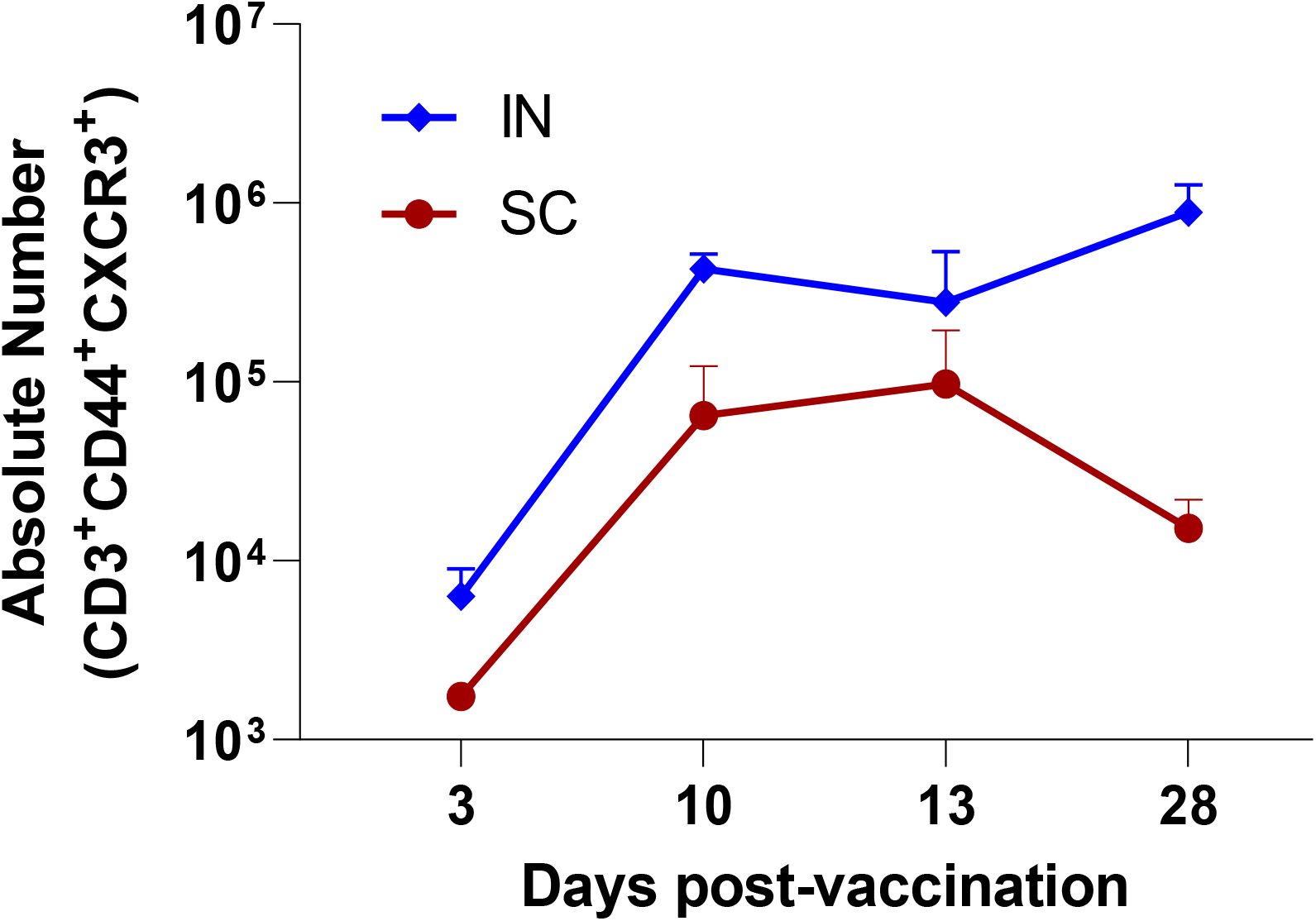
T cell recruitment to the lungs following BCG vaccination. Mice were vaccinated with BCG either via systemic (subcutaneous; SC) or mucosal (intranasal; IN) routes. On days 3, 10, 13, and 28 post-infection, mice were injected with fluorescently labeled anti-CD45 intravenously (i.v.) and then euthanized 3-5 minutes later, allowing sufficient time to stain cells in the vasculature but not those in the tissues. Lungs were extracted and digested with collagenase/DNase. Then, single cell suspensions were prepared for flow cytometric studies. Absolute numbers of CD3+CD44+CXCR3+ were calculated by multiplying the total number of viable cells recovered based on trypan blue staining and the percentage of the specific T cell subset detected by flow-cytometric analysis. The results demonstrate that on day 3 post-mucosal vaccination, T cells begin to infiltrate the lungs. The number of activated T cells recruited to the lung parenchyma decrease on day 28 in mice vaccinated with SC BCG.

### Protection by systemic BCG does not interfere with the anti-MAC effect of clarithromycin

Macrolides such as clarithromycin are one of the key groups of antibiotics used for treatment of pulmonary MAC. Macrolides are anti-inflammatory drugs [26] and it is important to study if the changes in MAC immunity induced by whole cell vaccines affect the ant-MAC activity of macrolides. Figure 5 shows that BCG does not interfere with anti-MAC activity of clarithromycin [provide overview of expt— suboptimal dug dose?).

**Figure 5.**
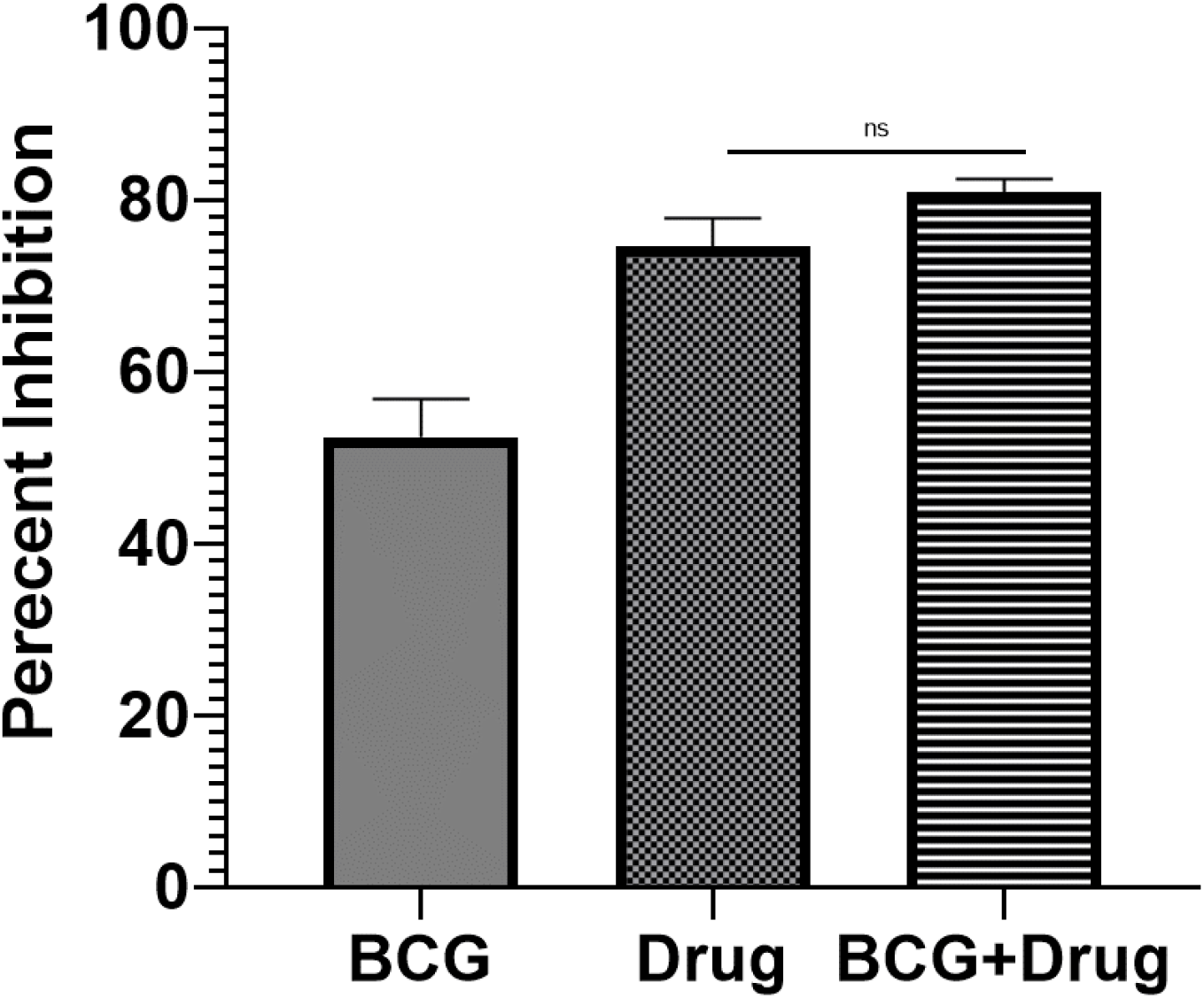
Effects of BCG vaccination on the anti-MAC effects of clarithromycin. Mice were vaccinated with BCG SC (1 × 10^7^ bacteria in 100 µL to base of tail). Four weeks after vaccination, mice were infected with aerosolized MAC at a concentration of 2 ×10^7^ CFU/ml. Two weeks after infection, some mice received clarithromycin at a concentration of 2 mg per 20 g via gavage 5 days a week. All mice were euthanized six weeks after infection, lungs homogenized and CFU quantified by culturing on 7H10 media. SC BCG did not interfere with anti-MAC activities of clarithromycin.

## Discussion

DAR-901 represents heat-killed NTM, *Mycobacterium obuense*, an investigational, whole-cell NTM vaccine derived from the Master Cell Bank of SRL172. The seed strain for SRL172 was initially identified as *M. vaccae* based on older phenotypic methods; more recently, 16s RNA sequence analysis has identified the organism as *M. obuense*, a rapid-growing NTM [21]. In a large NIH-sponsored Phase 3 trial in BCG-primed, human immunodeficiency virus (HIV)-infected patients in Tanzania, SRL172 was shown to be both safe and effective in the prevention of TB disease [27]. DAR-901 represents a scalable, broth-grown manufacturing method for agar-grown SRL172. DAR-901 has been shown to be safe in a Phase 1 trial that included HIV-negative and HIV-positive healthy adults with history of BCG vaccination [21] and in a Phase 2b trial among BCG-positive adolescents in Tanzania [25]. The results from multiple studies on SRL172 and DAR-901 indicate that this vaccine may not prevent Mtb infection but enhances Mtb immunity and prevents TB disease [25, 27]. Importantly, DAR-901 is itself an NTM and this study is the first study demonstrating that DAR-901 induces MAC cross-reactive T cell immunity and protects against an aerosol MAC challenge. The types of immunity providing protection against pulmonary MAC are not fully known. However, like Mtb, *M. avium* is an intracellular mycobacterium and requires Th1 T cells for effective control [23, 28, 29].

BCG, a live attenuated *M. bovis*, is part of childhood immunization programs in high TB endemic countries for protection against severe forms of TB. BCG is known to induce MAC cross-reactive immunity [23, 30]. In this study, we tested systemic and mucosal routes of BCG vaccination, and found that the mucosal route provided better protection against aerosol MAC challenges. Mucosal route of BCG vaccination has been shown to provide better TB protection compared to other routes of administration, presumably by eliciting unique transcriptomal molecular signatures [31, 32]. Our results showed that mucosal BCG vaccination led to recruitment of T cells to the lung much longer compared to systemic BCG vaccination and could be part of the reason why mucosal vaccination provides better MAC protection.

Different types of mice including BALB/c, C57BL/6, Beije, C3HeB/FeJ and nude strains have been used in trying to find the best small animal model for pulmonary *M. av*ium [22, 33, 34]. A study that compared BALB/c, C57BL/6, Beije and nude mice showed growth of *M. avium* in the lungs of all mice, but nude mice were found to be more susceptible to lung infection due to their deficient immune systems. BALB/c mice were found to be the best to study the impact of anti-mycobacterial treatment [22]. Therefore, in our study, BALB/c mice were used to determine the effects of vaccine-induced immunity on ant-MAC activity of clarithromycin. Macrolides including clarithromycin are known to have anti-inflammatory activities [35, 36]. The exact mechanisms determining how macrolides exert anti-inflammatory effects are not completely clear but it has been shown that macrolide treatment may lead to a decreased levels of TNF-γ and IL-12, cytokines which are important for Th1 immunity [36]. It is also not clear how the macrolide anti-inflammatory properties are related to anti-MAC activity or if interventions such as vaccinations which may lead to enhanced Th1 immunity affect the ant-MAC activity of macrolides. Our results clearly show that BCG vaccination does not interfere with anti-MAC activity of clarithromycin.

In conclusion, DAR-901 vaccination induces MAC cross-reactive immunity and protects against aerosol MAC challenge. In addition, mucosal BCG vaccination provides a better protection against aerosol MAC challenge compared to systemic BCG vaccination.

## Author Contributions

GA drafted the work, analyse results and prepare draft manuscript. KM and CC performed laboratory assays and enter data. CE supervised laboratory assays and reviewed manuscript.

## Data Availability Statement

The datasets for this study are available from the corresponding author upon request.

