## Supplementary Figure 1 for "Immunity against Mycobacterium avium induced by DAR-901 and BCG"

**Supplementary Results**

Supplement Figure 1.

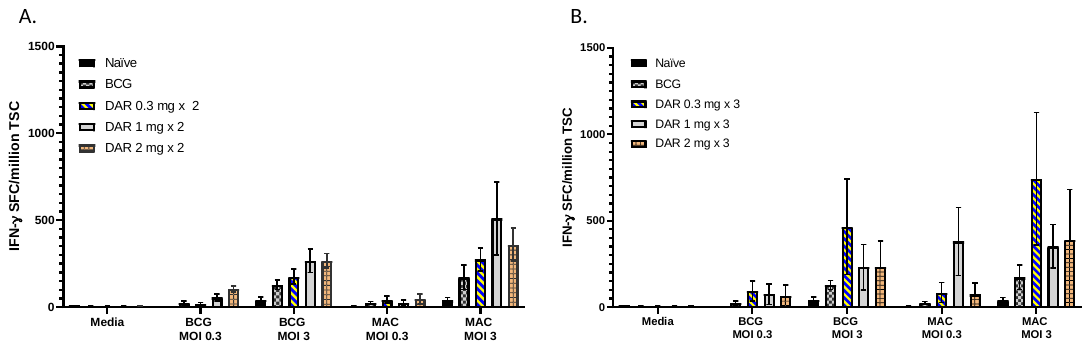

Suppl. Figure 1. *M. avium* cross reactive immunity induced by vaccination with BCG and DAR-901. Six-to 8-week-old female BALB/c mice were vaccinated with BCG (4 x 10^5^ intradermal) or different concentrations of DAR-901 (2 or 3 intradermal doses 2 weeks apart). Four weeks after last vaccination, mice were euthanized and splenocytes were used for IFN-γ ELISPOT assay. In the ELISPOT assay, splenocytes (5 x 10^5^ cells/well) were stimulated overnight with live BCG at multiplicity of infection (MOI) of 3, *M. avium* at MOI of 3, or media alone as a negative control. IFN-γ producing spots in each well were enumerated using a C.T.L. ImmunoSpot analyzer and software. The results are presented as spot forming cells (SFC, mean ± SE) per million total splenic cells (TSC). (A) shows results from mice vaccinated with BCG or 2 doses of DAR-901 a week apart at different concentrations. (B) shows results from mice vaccinated with BCG or 3 doses of DAR-901 a week apart at different concentrations.
